# Glio-SERS: Label-Free Molecular Profiling of Plasma Extracellular Vesicles in Brain Tumors Using SERS and Artificial Intelligence

**DOI:** 10.64898/2026.04.21.719934

**Authors:** Hulya Torun, Ugur Parlatan, Tim Valencony, Demir Akin, Chris Nguyen, Ozgur Albayrak, Furkan Kaysin, Ugur Aygun, Bharti Singal, Mehmet Ozgun Ozen, Rana Cansu Egitimci, Ibrahim Kulac, Oguz Baran, Goktug Akyoldas, Ihsan Solaroglu, Utkan Demirci

**Affiliations:** Bio-Acoustic MEMS in Medicine (BAMM) Lab, Canary Center at Stanford University, Department of Radiology, School of Medicine, Stanford University,{Citation} California, CA, 94304, United States; Biomedical Sciences and Engineering, Graduate School of Science and Engineering, Koç University, Istanbul, 34450, Turkiye; Koc University Research Center for Translational Medicine (KUTTAM), Koç University, Istanbul, 34450, Turkiye; Department of Electrical and Electronics Engineering, Koç University, Istanbul, 34450, Turkiye; Stanford Cryo-Electron Microscopy Center (cEMc), Stanford University, Palo Alto, CA, 94304; Koc University School of Medicine, Koc University, Sariyer, Istanbul, 34450, Turkiye; Department of Pathology, Koc University Hospital, Topkapi, Istanbul, 34010, Turkiye; Department of Neurosurgery, Koc University Hospital, Topkapi, Istanbul, 34010, Turkiye; Adjunct Research Professor, Loma Linda University School of Medicine, Basic Sciences, Physiology Division, Loma Linda, 92350, CA, USA; Department of Electrical Engineering (by courtesy), Stanford University, Stanford, CA, 94305, United States

**Keywords:** Glioblastoma, Extracellular vesicles, Liquid biopsy, Raman spectroscopy, Artificial intelligence

## Abstract

Extracellular vesicles are increasingly recognized as important carriers of disease-associated molecular information, yet robust methods for their isolation and molecular characterization from limited clinical samples remain challenging. Here, we present an integrated approach combining standardized EV isolation, label-free Surface-Enhanced Raman Spectroscopy (SERS), and artificial intelligence (AI) for comprehensive molecular profiling of small extracellular vesicles (sEVs) from human plasma. Here, we show systematically isolated and characterized plasma sEVs using ExoTIC in accordance with MISEV2023 guidelines, with SERS analysis revealing quantifiable spectral differences across samples from patients with glioblastoma (n=20) and meningioma (n=23) compared to healthy controls (n=30). Among the evaluated AI models, the convolutional neural network most effectively captured group-level spectral differences in sEVs, achieving accuracies up to 88% in this pilot cohort. Further, an EGFR-based spectral regression model was explored to examine molecular variability across sEV samples. Parallel proteomic analysis presented statistically significant differences in several proteins elevated in glioblastoma or meningioma. This label-free, rapid approach provides a proof-of-concept framework for sEV molecular profiling establishing the basis for broad validation studies across diverse diseases.

## Introduction

Brain tumors present unique challenges for molecular diagnosis^1^ and monitoring because repeated tissue sampling is often impractical and tumors frequently exhibit substantial spatial and temporal heterogeneity^2–4^. Although liquid biopsy approaches have transformed biomarker development in several systemic cancers^5–12^, translation to neuro-oncology has been slower due to the limited abundance of tumor-derived signals in peripheral blood and the biological complexity of central nervous system tumors^3,13–15^. This challenge is particularly relevant in glioblastoma (GB), where marked molecular heterogeneity further complicates biomarker development^3,14,16,17^.

Small extracellular vesicles (sEVs)^9,18–21^ have emerged as promising liquid biopsy analytes because they circulate stably in biofluids and carry proteins^20,22–25^, nucleic acids^20,21,26,27^, and other biomolecules reflective of their cells of origin^20,28–30^. sEVs have shown potential for non-invasive disease profiling^31–35^, treatment response monitoring^32,36^, molecularly informed patient stratification^32,37^ across multiple clinical contexts, including oncology^5–11^, cardiovascular^38–41^, and neurological^6,42–47^ diseases. Their cargo is particularly attractive as a source of disease-relevant molecular information, due to their abundance, stability, and capacity to reflect disease-specific biological processes^5,27,30^. However, robust approaches for isolating and molecularly profiling circulating sEVs from plasma remain an important challenge, particularly when attempting to detect tumor-associated molecular signals in brain tumor patients ^48–54^.

Label-free molecular profiling approaches offer a complementary strategy for analyzing complex EV populations without relying on predefined biomarker panels. Surface-enhanced Raman spectroscopy (SERS), an ultra-sensitive variant of Raman spectroscopy, enables detection of molecular vibrational signatures and provides biochemical fingerprints of biological samples^14,55–57^. When combined with artificial intelligence (AI)-based molecular pattern recognition, SERS has demonstrated potential for detecting disease-associated molecular patterns in complex biofluids^14,57^, including blood^58,59^, urine^60,61^ saliva^62^, cell culture^57,63,64^, cerebrospinal fluid (CSF)^14,65,66^, and tumor tissue ^67,68^. Nevertheless, integration of standardized EV isolation, label-free spectroscopic profiling, and AI-based analysis for plasma samples from brain tumor patients remains relatively underexplored.

Here, we present Glio-SERS, an integrated framework (**Fig. 1**) combining standardized extracellular vesicle isolation using the Exosome Total Isolation Chip (ExoTIC)^27^ (**Fig. 1b**), label-free SERS molecular fingerprinting (**Fig. 1c-e**), and AI-based classification for disease-associated molecular stratification of plasma sEVs (**Fig. 1c-f**). Using plasma samples from 73 individuals, including patients with primary glioblastoma, (GB, n=20), meningioma (MNG, n=23), and healthy controls (HC, n=30), we evaluated a two-stage proof-of-concept approach: first, detection of a brain tumor-associated molecular signal, and second, exploratory tumor-type stratification. Proteomic analyses (**Fig. 1c**) were performed in parallel with the SERS analyses, providing independent molecular context for interpreting AI-derived spectral patterns. Together, this study evaluates the feasibility of integrating standardized EV isolation, label-free spectroscopy, and AI-enabled analysis for minimally invasive molecular profiling of circulating sEVs in brain tumors.

**Figure 1.**
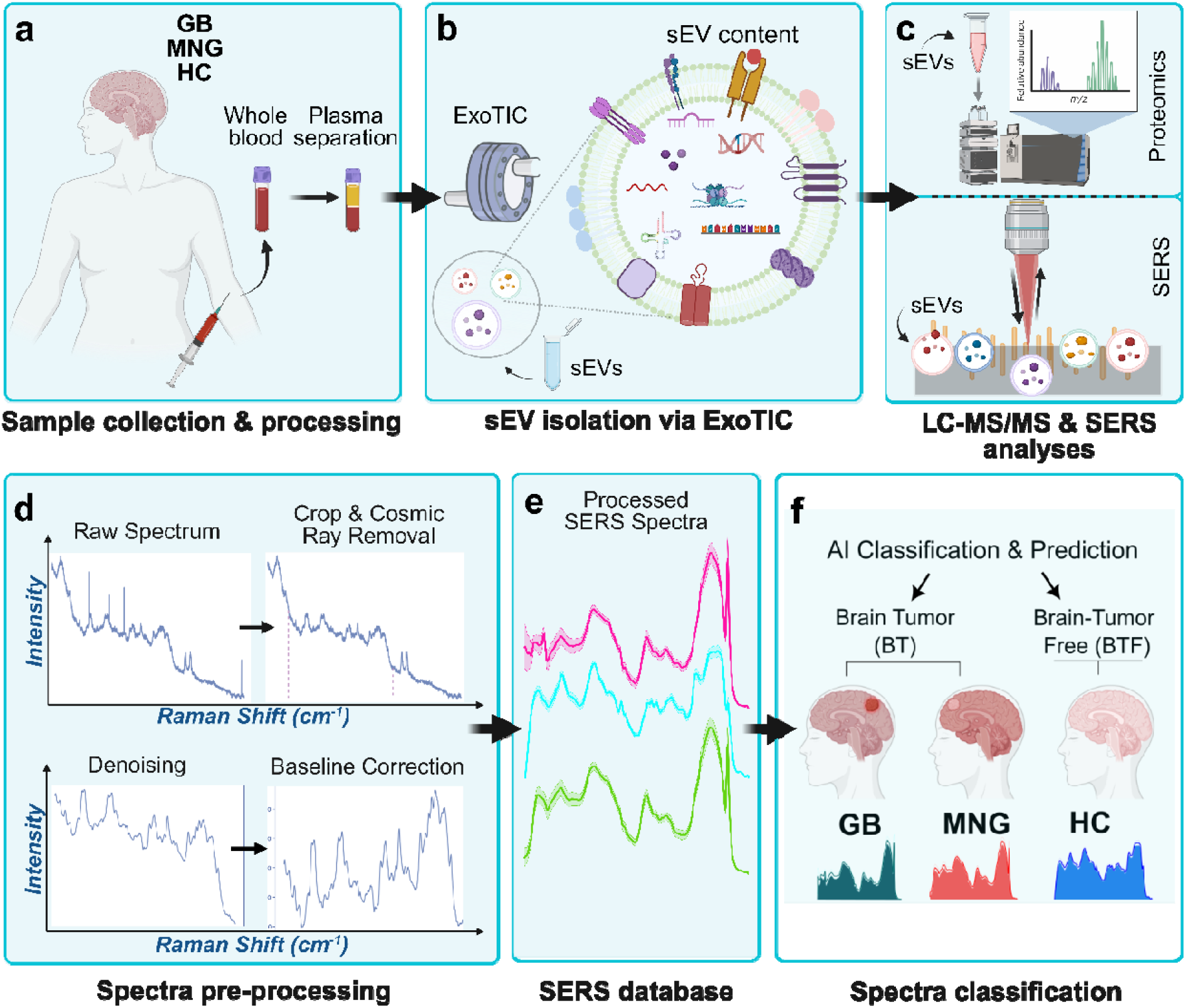
Integrated ExoTIC-SERS-AI based sEV profiling approach enables characterization of plasma-derived small extracellular vesicles (sEVs) from brain tumor patients (Glioblastoma and Meningioma) and healthy controls (HC). This study combines plasma collection, small extracellular vesicle (sEV) isolation, surface-enhanced Raman spectroscopy (SERS), and artificial intelligence (AI) to develop a non-invasive analysis framework for distinguishing glioblastoma (GB), meningioma (MNG), and healthy control (HC) samples. **a**, Blood samples were collected from participants with and without brain tumors, followed by plasma separation. **b**, Plasma sEVs were isolated using the Exosome Total Isolation Chip (ExoTIC) and characterized according to MISEV2023 guidelines.**c**, sEVs were analyzed on gold-patterned substrates using SERS to capture molecular fingerprints. **d**, Spectral data were pre-processed through cosmic ray removal, denoising, and baseline correction. **e**, The curated spectral dataset was used for training and testing AI classification models. **f**, Multiple AI algorithms were applied to classify sEV SERS spectra from GB, MNG, and HC groups. Created in BioRender. Torun, H. (2026) https://BioRender.com/j64p037.

## Materials and Methods

### Sample Collection and Inclusion Criteria

Blood samples were collected under the approval of the Koç University Institutional Review Board (#2019.328.IRB2.106) with the oral and written consent of the patients and/or their legal guardians.

Blood samples from IDH-wild type primary GB (WHO grade IV) and MNG (WHO grade I and II) patients were collected in EDTA tubes prior to tumor resection in the operation room under anesthesia. Recurrent cases and/or cases with prior radiotherapy were excluded from the study. Diagnoses were performed according to WHO 2021 CNS Tumor classification guidelines and standard operating procedures by the Koç University Department of Pathology through the evaluation of tumor samples obtained from the patients. Blood samples from healthy individuals were collected from volunteers without any known tumors or other malignancies as well as any other chronic diseases.

### Blood Processing and sEV Isolation

Blood samples were centrifuged at 2000 rpm and 4°C for 20 minutes, after which the upper layer of the supernatant (plasma) was carefully collected without disturbing the buffy coat. Plasma was aliquoted and stored at -80°C for future use. Before sEV isolation, plasma samples were thawed. A volume of 500 µl of plasma was used for sEV isolation using the Exosome Total Isolation Chip (ExoTIC)^27^ as a well-established tool for various applications, sample types and disease applications^27,57,69,70^ The ExoTIC membranes were incubated in PBS, collected from the ExoTIC collection chamber, at 4°C overnight to facilitate sEV release. ExoTIC system isolated sEV and particles that range from 30-220nm (**Fig 1b**). sEVs suspended in phosphate-buffered saline (PBS, Cytiva) were collected the following day from the ExoTIC membranes and stored at -80°C until further use.

### sEV Characterization

sEVs were characterized using Transmission Electron Microscopy (RT-TEM), Cryo-Electron Microscopy (CryoEM), Interferometric Scattering Microscopy (iSCAT), Nanoparticle Tracking Analysis (NTA), Bead-Captured Flow Cytometry (BC-FC), and Western Blotting (WB).

#### TEM Analysis of sEVs

sEVs were diluted to the optimal concentration (10x–100x) and subjected to negative staining. A 10 µl aliquot of the sEV suspension was pipetted onto a glow-discharged grid (Electron Microscopy Sciences, FCF-300-CU) and allowed to settle for 3 minutes. Subsequently, three drops of 1% uranyl acetate were applied sequentially, with each drop left on the grid for 1 minute. Excess uranyl acetate was carefully removed using filter paper. To remove residual uranyl acetate, three drops of distilled water were applied to the grid, and the excess water was similarly blotted with filter paper. The grid was then air-dried for approximately 10 minutes before the imaging. sEV imaging was performed at 100 kV using a JEOL JEM 1400 120 kV TEM microscope (JEOL USA, Inc.), equipped with a Gatan UltraScan digital high-resolution camera (Pleasanton, CA) (**Fig. 2a**).

**Figure 2.**
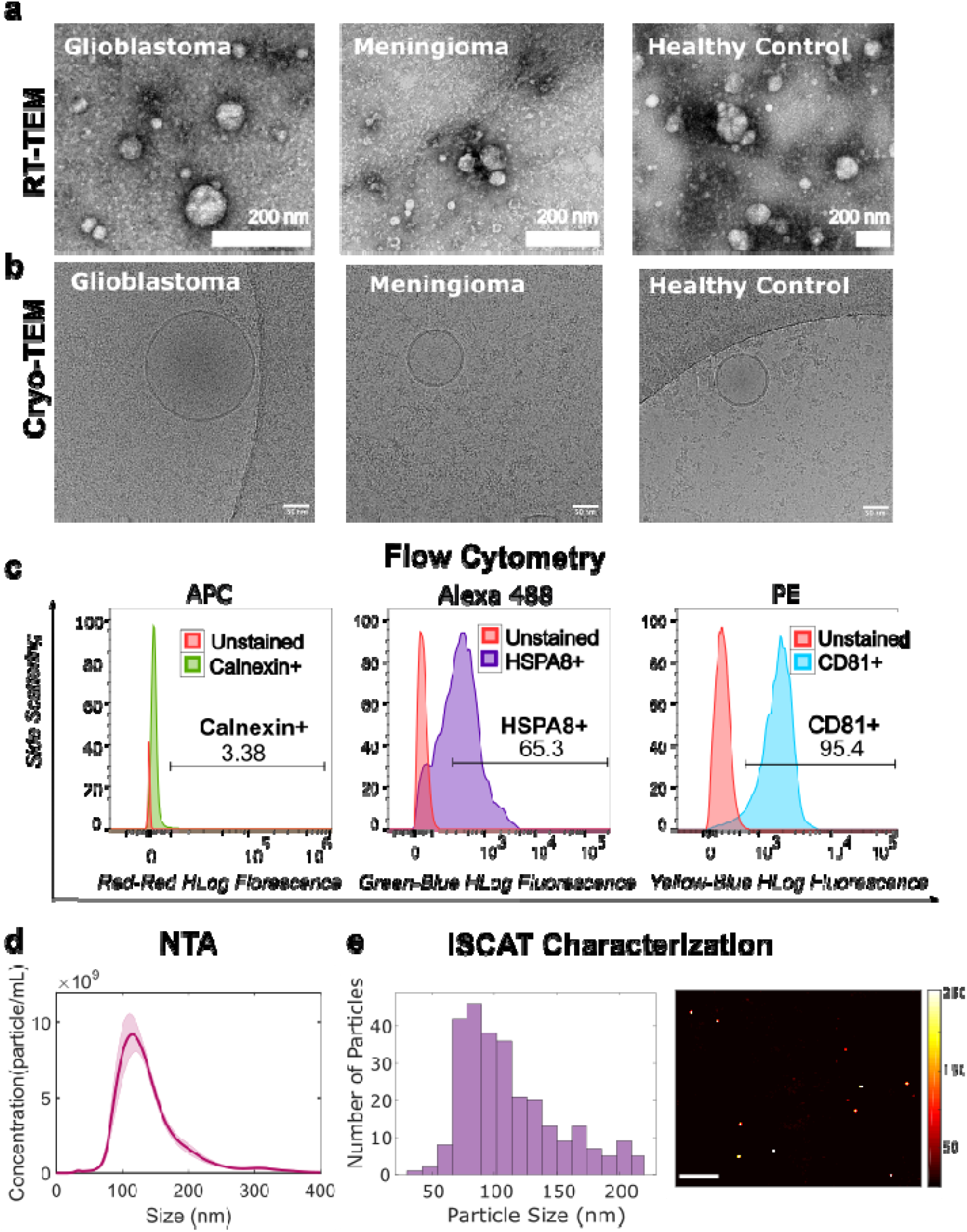
Multimodal characterization confirms the integrity, size distribution, and surface profiles of plasma-derived extracellular vesicles. A combination of high-resolution imaging and particle analysis techniques verified the structural integrity, size heterogeneity, and molecular features of isolated small extracellular vesicles (sEVs). **a**, Room-temperature transmission electron microscopy (RT-TEM) reveals the typical spherical morphology of sEVs (scale bar: 200 nm). **b**, Cryogenic transmission electron microscopy (CryoEM) visualizes the native ultrastructure of intact vesicles under vitrified conditions (scale bar: 50 nm). **c**, Bead-capture flow cytometry (BC-FC) shows surface marker expression on sEVs, confirming vesicular identity. (Staining presented as percentage). **d**, Nanoparticle tracking analysis (NTA) provides particle size distribution and concentration in solutions. **e**, Interferometric scattering microscopy (iSCAT) enables label-free imaging and precise size measurement of individual sEVs at the single-particle level.

#### Cryo-EM Analysis of sEVs

High-resolution imaging of isolated sEVs was performed using cryo-electron microscopy (Cryo-EM). Quantifoil Holey Carbon 1.2/1.3 Cu 300 mesh grids were glow-discharged using the PELCO easiGlow™ (TED PELLA Inc.) for 45 seconds at a current of 15 mA. A 3 μL of the sEV sample was applied to the glow-discharged grids and blotted using a Vitrobot Mark IV (FEI) for 3 seconds at a blot force of 3, with a 10-second wait time, at a temperature of 22°C and 100% humidity.

The grids were then plunge-frozen in liquid ethane without drain time, clipped, and loaded into a ThermoFisher Scientific Glacios™ Transmission Electron Microscope (Cryo-TEM) operating at 200 kV. The specimens were imaged at 120,000x magnification with a pixel size of 1.2 Å, using The ThermoFisher Scientific Falcon4i Direct Electron Detector and EPU software (ThermoFisher Scientific). The total accumulated electron dose did not exceed 60 e□/Å^2^, and the defocus range was set at -3 microns with a dose rate of 6.99 e□/pix/sec. (**Fig. 2b**).

#### NTA Analysis of sEVs

sEV size and concentration distribution were evaluated using nanoparticle tracking analysis (NTA) on a Nanosight NS300 (Malvern Panalytical, UK) equipped with NTA software version 4.2. A 532 nm green laser (50 mW) was used, with the camera level set between 8 and 10 and a detection threshold of 5. sEV samples were diluted to an optimal concentration (100x–1000x) to achieve a particle count of 20–100 particles per frame. NTA analyses were then performed (**Fig. 2d**).

#### Flow Cytometry Analysis of sEVs

sEVs were incubated with aldehyde sulfate latex beads (ThermoFisher) for 15 minutes at room temperature (RT) on a roller. The sEV-bead complexes were then blocked using 2% BSA for 2 hours, followed by a 30-minute incubation with 100 mM glycine to reduce nonspecific binding. The complexes were washed with cold PBS to remove unbound sEVs and beads from the solution.

The sEV-bead complexes were incubated overnight with primary antibodies, including anti-Calnexin (#PA1-30197, Invitrogen), anti-CD81 (#SAB4700232, Sigma-Aldrich), and anti-HSPA8 (#NB100-41377, Novus Biologicals). The following day, the samples were centrifuged and washed with cold PBS to remove unbound antibodies. Secondary antibodies (#A28175, #A-10931, and #31860, Invitrogen) were then added, and the samples were incubated for 2 hours. After centrifugation and washing with cold PBS, the samples were analyzed by flow cytometry using a Guava easyCyte flow cytometer, and data were processed with FlowJo v10 software (**Fig. 2c**).

Control samples, including only sEVs, only beads, and beads with sEVs, were used to identify the purest sEV population, as recommended by MISEV2023 guidelines^30^. To isolate the sEV population, Calnexin-negative (Calnexin□) sEVs were gated first. A multi-gating strategy was then applied to identify CD81-positive (CD81□) and HSPA8-positive (HSPA8□) populations, as well as double-positive and double-negative populations for these markers (**Fig. 2c**).

#### Interferometric Scattering Microscope Analysis of sEVs

We developed a custom interferometric scattering microscope specifically designed for the quality control of isolated sEVs^71^. The microscope operates on a thin-film substrate composed of SiO□/Si to capture high-resolution interferometric images. Illumination is provided by an LED (Thorlabs M530L4, 530 nm) focused on the back focal plane of the objective lens (Nikon LMPlanFL, 50×) to ensure homogeneous illumination. Scattered light from the sEVs, combined with reflected light from the substrate, is captured by the objective lens and transmitted to a camera to generate interferometric images. A motorized stage (Standa) is employed to facilitate fine focus adjustments.

For imaging, sEV samples were deposited onto a substrate consisting of a 100 nm SiO□ thin film layer atop a Si base and allowed to dry for 15 minutes. iSCAT images of the sEVs were acquired over a large field of view (200 µm × 200 µm), enabling sensitivity to individual particles. Each particle within the field of view was detected using a custom MATLAB script, and the size distribution of the particles was calculated, as shown in **Figure 2e**.

#### Western Blot Analysis of sEVs

Western blot analysis of sEVs was conducted following a previously described protocol. The analysis used Anti-Calnexin (#PA1-30197, Invitrogen,Carlsbad, CA,United States), Flotillin1 (#A6220, Abclonal,Woburn, MA, USA), and Anti-CD63 ((#556019, BD Pharmingen,San Jose, CA, USA) antibodies in accordance with the MISEV2023 guidelines^30^. (**Fig. 2c and Supplementary Fig. 2-3**)

### Surface-Enhanced Raman Spectroscopy Analysis of sEVs

#### Experimental Setup

A customized Raman spectroscopy system was developed to provide enhanced flexibility for multi-modal and multi-functional analysis while maintaining sensitivity comparable to state-of-the-art instruments. A diode laser (CrystaLaser, 785 nm, maximum output power: 130 mW) was used, with the output power adjusted via a half-wave plate and a linear polarizer to ensure optimal performance without damaging the sensitive gold (Au) substrate.

The adjusted laser beam was directed to the sample plane of the microscope through a long-pass dichroic mirror (Thorlabs, DMLP 805) and a microscope objective (10x, NA 0.22). A motorized XYZ stage (Standa) was used to facilitate lateral scanning of the sample surface and precise focal plane adjustments. The scattered photons were collected using the same objective lens in a 180-degree backscattering geometry. Elastic (Rayleigh) scattering was filtered out by the dichroic mirror, while inelastic (Stokes) scattered photons were directed to a multimode fiber via an ultra-steep long-pass filter (Semrock) and a fiber collimator (Thorlabs, F220SMA-780). The collected signal was then transmitted to a Raman spectrometer (StellarNet, HyperNova) equipped with a CCD camera (Andor iVAC 316), which was cooled to -60°C to minimize noise and enhance sensitivity.

#### Sample Preparation

For SERS measurements, hydrophobic Au-coated surfaces (Silmeco ApS, Copenhagen, Denmark) were utilized to optimize signal enhancement. A 2 µL aliquot of sEV sample was deposited onto the Au surface and allowed to dry for 15 minutes. Each SERS surface was used only once to ensure consistent enhancement performance.

#### Data Collection

A connection between the PC and the CCD spectrometer was established using the Andor MATLAB R2024A SDK. A custom MATLAB script was developed to automate surface scanning with predefined parameters. The laser output power was fine-tuned to 30 mW (approximately 20 mW at the sample plane). Real-time spectral feedback was employed to identify the optimal focal plane for data acquisition.

Once the focus was optimized, spectral acquisition was performed with the following parameters: an exposure time of 1 second, a scanning step size of 2.5 µm, and 21 steps along each axis (X and Y). This configuration yielded a total of 441 spectra per sample, ensuring robust and comprehensive spectral data for analysis.

#### Dataset Preprocessing and Artificial Intelligence Models

A total of 32,193 spectra were collected from three classes: GB, MNG, and HC. The dataset was preprocessed using a series of steps implemented with the RamanSPy^72^ library in Python. First, the regions outside the fingerprint region were cropped from each raw spectrum. Cosmic rays were removed using the Whitaker-Hayes algorithm, and spectra were smoothed using the Savitzky-Golay algorithm^73^. Fluorescence background was leveled and removed using the Improved Asymmetric Least Squares (IALS) baseline correction algorithm. Finally, each spectrum was normalized to its intensity, facilitating easier comparison of Raman profile shifts (**Fig. 3a**).

**Figure 3.**
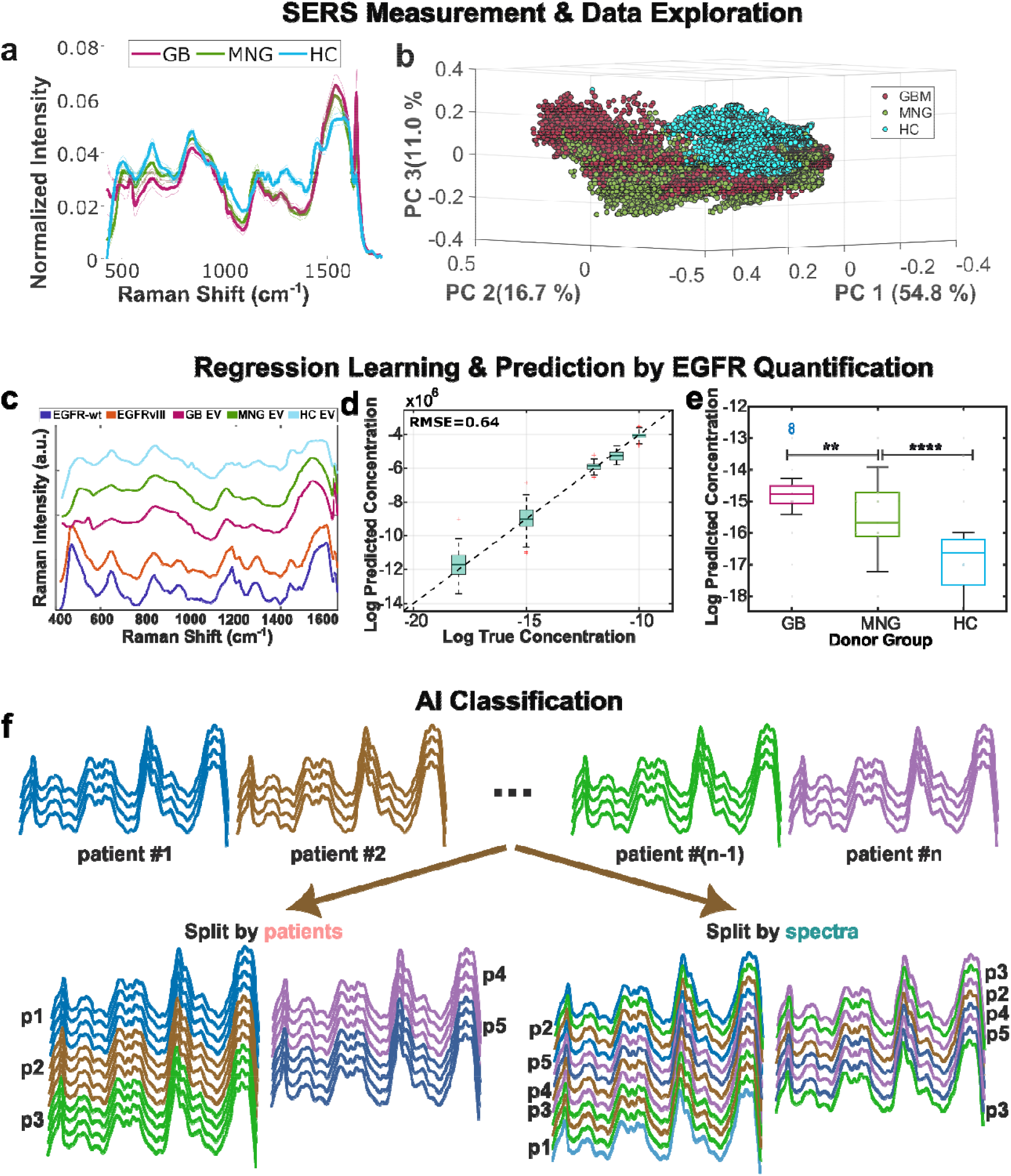
SERS-based profiling of plasma-derived extracellular vesicles enables detection of EGFR signatures and molecular classification of brain tumors. Surface-enhanced Raman spectroscopy (SERS) was applied to small extracellular vesicles (sEVs) isolated from plasma samples to detect EGFR signatures and discriminate between glioblastoma (GB), meningioma (MNG), and healthy control (HC) participants. (p < 0.05, ** p < 0.01, *** p < 0.001, **** p < 0.0001) **a**, Representative SERS spectra of sEVs from individuals with GB, MNG, and HC, highlighting spectral differences across groups. **b**, Three-dimensional principal component analysis (PCA) showing group-level clustering of SERS spectra. **c**, SERS spectra of purified EGFR wild type (wt), EGFRvIII, and sEVs from GB, MNG, and HC samples. Green-shaded regions mark spectral shifts characteristic of EGFR signatures observed in clinical samples. **d**, Regression model trained to predict EGFR (wt) concentrations from SERS spectra. Predicted concentrations from the 20% hold-out test set are plotted against true values; root-mean-square error (RMSE) = 0.64. **e**, Model-predicted EGFR (wt) concentrations for sEVs from GB, MNG, and HC samples. Box plots display inter-group differences (p < 0.01, ** for GB vs MNG; p < 0.0001, **** for all other comparisons; two-sided Mann–Whitney U test). **f**, Schematic of the artificial intelligence-based pipeline for SERS data analysis. Two data-splitting strategies were used: patient-wise (left), where each participant’s 441 spectra (p1, p2, …, pn) were treated as a unit; and spectrum-wise (right), where spectra were randomly split across participants. Each participant contributed 441 SERS spectra; training and test sets were generated based on predefined split ratios.

After preprocessing, the data distribution was analyzed (**Supplementary Fig. 5a-d**) using t-Distributed Stochastic Neighbor Embedding (t-SNE), a non-linear dimensionality reduction technique^74^. This method embeds high-dimensional data into two- or three-dimensional space, allowing visualization of class distribution and potential similarities. Similar spectra formed clusters in the reduced-dimensional space, while dissimilar spectra were more dispersed, providing insights into class-specific patterns.

For patient state prediction, we compared the performance of six pre-defined models: Convolutional Neural Network (CNN)^75^, Neural Network (NN)^76^, Random Forest Classifier^77^, Support Vector Classifier (SVC)^78^, Quadratic Discriminant Analysis (QDA)^79^, and XGBoost^80^. These models were evaluated to determine their accuracy and reliability in predicting patient states (**Supplementary Fig. 7-8, Table 1, Supplementary Table 1**).

**Table 1.** AI classification results for brain tumor (BT: GB and MNG) vs healthy control (HC) groups. The classification results are presented for 80:20 train-test split ratio Area Under the Curve (AUC), Specificity, Sensitivity, and Weighted f1 Score values are presented for 80:20 patient-wise train-test split ratio. (CNN: Convolutional Neural Networks, NN: Neural Networks, RF: Random Forest, SVM: Support Vector Machine, QDA: Quadratic Discriminant Analysis)

| Models/<br>Train-Test<br>Split Ratio | AUC<br>80:20 | Accuracy | Specificity (BT<br>vs HC)<br>for 80:20 | Sensitivity (BT<br>vs HC)<br>for 80:20 | Weighted f1<br>Score<br>for 80:20 |
| --- | --- | --- | --- | --- | --- |
| **CNN** | 0.94 | **0.88** | 0.94 | 0.85 | 0.87 |
| **NN** | 0.78 | **0.5** | 0.77 | 0.53 | 0.43 |
| **RF** | 0.95 | **0.79** | 0.91 | 0.79 | 0.77 |
| **SVM** | 0.97 | **0.79** | 0.91 | 0.8 | 0.78 |
| **XGBoost** | 0.94 | **0.76** | 0.9 | 0.76 | 0.75 |
| **QDA** | 0.84 | **0.63** | 0.82 | 0.62 | 0.62 |

#### Statistical Analysis

A Kolmogorov-Smirnov test was conducted on selected intense Raman peaks at 456, 587, 715, 761, 919, 1069, 1219, 1254, 1304, and 1422 cm□^1^, which revealed a non-normal distribution of the data. Consequently, non-parametric statistical methods were applied. To evaluate differences among the three groups in our study (GB, MNG, and HC), a Kruskal-Wallis test was performed on the selected peaks. The analysis showed significant differences in Raman intensity distributions among these groups (*p* < 0.001). The distributions of spectral data for the selected peaks are visualized using violin plots (**Supplementary Fig. 4b-d**).

#### Regression analysis

To assess the presence of amino acids in EGFR protein samples, we first identified the most and least frequent amino acids. A dataset was then constructed, incorporating Raman peak intensities from different concentrations of EGFR (#10001, Sino Biological Inc.) and EGFRvIII (#29662, Sino Biological Inc.) protein preparations and their constituent concentrations (v:v ratios). This dataset was used to develop a linear regression model. The data was split into training (80%) and testing (20%) sets, and k-fold cross-validation (k=5) was employed to validate the model (**Fig. 3d**).

Using this model, we validated the presence of selected amino acids in the SERS spectra of EVs by analyzing their most dominant Raman peaks as reported in the literature. The performance of the model was assessed by calculating the root mean square error (RMSE) values for the prediction.

### LC/MS analysis of plasma sEVs

sEVs were isolated using 50 mM ammonium bicarbonate for proteomics analysis. sEV proteins were exposed by disrupting the lipid bilayer with 1.5% sodium dodecyl sulfate (SDS) before further processing. Cold acetone was used to remove SDS from the samples. BCA assay was used for protein quantification. 25 µg of protein was used per sample. All analyses were performed with three technical replicates.

1.5 µg of tryptic peptides were loaded onto an Acclaim PepMap C18 trap column (Thermo Fisher Scientific) coupled to a Dionex Ultimate Rapid Separation Liquid Chromatography HPLC system (Thermo Fisher Scientific) at a flow rate of 5 µL/min for 10 minutes. Separation of tryptic peptides was achieved using reversed-phase chromatography on a 25 cm-long C18 analytical column (New Objective) packed in-house with ReproSil-Pur 120 C18 AQ resin (Dr. Maisch GmbH). Peptides were eluted and ionized via a Nanospray Flex ion source (Thermo Fisher Scientific) with a 1.8 kV voltage and analyzed using an LTQ-Orbitrap Elite mass spectrometer (Thermo Fisher Scientific).

The chromatography gradient was programmed as follows: a flow rate of 0.4 µL/min was maintained throughout. Mobile phase A (0.1% formic acid in water) and mobile phase B (0.1% formic acid in acetonitrile) were used, starting at 98% A and 2% B for 10 minutes. This was followed by a gradual increase to 35% B over 100 minutes, then to 85% B over 2 minutes, with a 7-minute hold. Re-equilibration of the analytical column was performed before each subsequent injection. Each sample was analyzed in triplicate.

The top 10 most abundant ions from each MS1 scan were selected for higher-energy collision-induced dissociation (HCD) at 35 eV in a data-dependent acquisition mode. MS1 scans were performed with a resolution of 60,000, an FT AGC target of 1e^6^, and an m/z scan range of 400–1800. MS2 scans were acquired with an AGC target of 3e^4^, and dynamic exclusion was enabled for 30 seconds.

### Proteomics data analysis

Raw data files from each LC-MS run were analyzed using Byonic 4.0.12 (Protein Metrics, San Carlos, CA). Data were searched against the Swiss-Prot database, incorporating the 2024 reference human proteome with 20,692 protein entries. Search parameters specified trypsin as the digestion enzyme, allowing up to two missed cleavages. A precursor mass tolerance of 10 ppm and a fragment mass tolerance of 0.5 Da were applied. Fixed modifications included carbamidomethylation of cysteine, while variable modifications accounted for methionine oxidation and asparagine deamination.

To ensure high-confidence peptide identification, spectra with a false discovery rate (FDR) greater than 1% were excluded. All experimental conditions were performed in triplicate, with three technical replicates per condition. Protein quantification was based on the specific signal intensity of each protein, normalized to the average signal across all samples, enabling relative abundance determination within the dataset.

Normalization procedures were applied to adjust relative abundance values to a standard normal distribution with a mean of 0 and a standard deviation of 1, ensuring comparability across samples. Supervised clustering was then conducted using Pearson correlation across all biological conditions, including permutations. Only clusters showing significant correlations (*p* < 0.01) were retained for further analysis. Significantly enriched clusters were analyzed using overrepresentation analysis via the WebGestalt tool (http://www.webgestalt.org/). This analysis identified enriched pathways and biological processes, providing insight into the functional implications of proteomic data.

## Results

### Characterization of sEVs

Plasma-derived sEVs were characterized using complementary imaging and molecular techniques to assess size distribution, morphology, and marker expression in accordance with MISEV2023 guidelines^21^. Room temperature transmission electron microscopy (RT-TEM), cryogenic TEM (CryoEM) and interferometric scattering microscopy (iSCAT) revealed that the majority of sEVs were spherical and within the 90-150 nm range (**Fig. 2a,b,e)**, and **Supplementary Fig. 1b**), consistent with size distributions measured by nanoparticle tracking analysis (NTA) results (**Fig. 2d, Supplementary Fig. 1a**). Bead-Assisted Flow Cytometry (BC-FC) analysis resulted in 93% of particles were stained for Calnexin (-), 83.4% of particles were stained for Calnexin (-) and CD81 (+), 80.7% of particles were stained for Calnexin (-) and HSPA8 (+), and 68.9% were stained for Calnexin (-), CD81 (+), and HSPA8 (+) antibodies (**Fig. 2c, Supplementary Fig. 2)**. The WB analysis on sEVs using CD63, TSG101, Flot1, showed protein bands at 50, 47, and 50 kDa, respectively (**Supplementary Fig. 1c**). Calnexin, a cellular protein control, band was not observed. These characterization results demonstrated sEV purity and quality, in alignment with the MISEV2023 guidelines^30^.

### Surface-Enhanced Raman Spectroscopy (SERS) Data Analysis of sEVs

To analyze group-specific variations in SERS spectra of sEV measurements, we first compared the average SERS spectra of the GB, MNG, and HC groups (**Fig. 3a**). Following pre-processing (baseline correction, smoothing, normalization) as detailed in Methods (see: Dataset Preprocessing and Artificial Intelligence Models), we applied Principal Component Analysis (PCA) on the normalized dataset. The 3D visualizations of PCA scores plot (**Fig. 3b** and **Supplementary Video 1**) demonstrate clustering of samples, with the first three principal components (PC 1-3) explaining 54.8%, 16.7%, and 11% of the variance, respectively. T-SNE distribution of all donor groups were also shown (**Supplementary Fig. 4**).

To explore the spectral features contributing to group differentiation, we analyzed the PCA loading plot of the dataset for the first three principal components (**Supplementary Fig. 5a**). Next, we examined the intensities at three key Raman peaks (1069 cm^-1^, 1304 cm^-1^, and 1422 cm^-1^) using violin plots (**Supplementary Fig. 5b-d**). These plots provided significant group-wise differences between GB, MNG, and HC, especially at 1422 cm^-1^ (GB *vs*. HC: p<0.05; MNG *vs*. HC: **p<0.01; GB *vs*. MNG: ***p<0.001). Distinct spectral shifts at these Raman peaks correspond to biomolecular fingerprint differences including proteins, and lipid alterations, highlighting molecular heterogeneity between tumor types.

### Regression-Based Evaluation of EGFR-Related Spectral Differences in Plasma sEVs Reveals Molecular Variations Across Sample Groups

Epidermal growth factor receptor (EGFR) is one of the most frequently altered oncogenes in IDH-wildtype GB, with amplifications, mutations, or splice variants in ∼60% of tumors, (amplification alone in ∼40%) ^16,17,81,82^. These alterations drive tumor growth, progression, and resistance. Secondary alterations, such as constitutively active EGFRvIII variant, further enhance oncogenic signaling^83^. EGFR and its oncogenic variants, including EGFRvIII, are consistently found in sEVs from GB cells and patient biofluids, reflecting tumor molecular status and enabling minimally invasive monitoring^84–87^. To exploit these molecular signatures, we used SERS to profile sEVs and the spectral differences between purified EGFR wild-type and EGFR mutant proteins (**Fig. 3c**) via developing a regression learning model (**Fig. 3d**). The multivariable parameter set extracted from the EGFR SERS analysis was then applied to the sEV spectra. **Fig. 3e** shows the model outputs as box plots that show the distributions of the predictions per each group, supporting the possibility that SERS-based sEV profiling may capture molecular patterns associated with tumor presence for GB^84,88^. **Supplementary Fig. 6** shows the outputs of the same approach when the spectra are scanned for the most predictive wavenumbers.

To develop a quantitative prediction model, we first compared the average spectra corresponding to EGFR-wild type protein (wt), EGFRvIII protein, plasma sEVs from GB and HC patients to identify the SERS intensity differences as in **Fig. 3c**. Then, we built a linear regression learning model using the whole EGFR-wt spectra. The predictive capability of this model is illustrated in **Fig. 3d**. Our regression model performance (with RMSE = 0.64) suggests that the model can approximate EGFR-associated spectral variation within this experimental setting. Finally, the model was applied on the sEV measurements across three donor groups. Statistically significant differences were found between GB-derived sEVs and those from other groups, as demonstrated in **Fig 3e**. This study covers use of an EGFR regression model to reveal spectral shifts between GB, MNG and HC groups in this pilot dataset. There are many proteins in plasma and plasma EVs contributing to the EV-SERS spectra. Rather than measuring EGFR levels, **Fig. 3e** showcases how our linear model effectively separates the clinical groups. We further investigated the relation between the regression model prediction and the EGFR copy number information obtained from molecular pathology analysis (using Next Generation Sequencing-NGS) data for four EGFR amplified samples (**Supplementary Table 1**).

### AI-Enabled Spectral Profiling of Plasma sEVs Reveals Group-Specific Molecular Signatures

To evaluate the predictive potential of plasma-derived sEVs profiled by SERS, we applied machine learning models to classify patients into HC, MNG, and GB groups. The analysis followed a two-stage classification strategy. First, we grouped MNG and GB cases under a unified brain tumor (BT) label and assessed whether sEV spectra could distinguish individuals with brain tumors from healthy individuals (**Table 1**). Second, we performed a more refined classification task to differentiate between MNG and GB subtypes.

Two approaches were employed to partition the spectral dataset into training and testing sets: splitting by individual spectra^89^ and splitting by patient^90^ (**Figure 3f**). The former method, where individual spectra from the same patient may appear in both training and testing sets, yielded high classification accuracies ranging from 80.3% to 99.7% across all models for the BT vs HC task (**Supplementary Fig. 8, Supplementary Table 2)**). However, this approach may artificially enhance performance by allowing the model to learn patient-specific features. In contrast, patient-wise splitting, where all 441 spectra from a single individual were allocated exclusively to either the training or the testing set, better reflects clinical deployment scenarios and reduces overfitting risks (**Fig. 3f**). Using this more stringent approach, six classification models - convolutional neural networks (CNN) (**Supplementary Fig. 7**), neural networks (NN), random forest (RF), support vector machines (SVM), XGBoost, and quadratic discriminant analysis (QDA) - were evaluated across three train-test split ratios including 80:20 (**Table 1**), 60:40 and 40:60 (**Supplementary Fig. 9, Supplementary Table 2**), with classification performance reported for both tasks.

In this exploratory analysis within the BT *vs*. HC dataset (**Table 1, Supplementary Fig. 9a-d)**, CNN achieved the highest classification performance (87.6%), specificity (93.8%), sensitivity (85.0%), and a weighted F1 score of 87.0%, when trained on an 80:20 split. RF and SVM models followed with accuracy values of 78.9% and 79.3%, respectively, while XGBoost and QDA reached 76.3% and 63.5%. The NN model underperformed with 50.0% accuracy. As the proportion of training data decreased, performance dropped across all models, although CNN retained relatively high accuracy at 73.3% (60:40 split) and 66.6% (40:60 split), outperforming other classifiers under limited training conditions (**Supplementary Table 2** and **Supplementary Fig. 9a-d**).

In the GB *vs*. MNG classification task **(Supplementary Fig. 9e-h)**, CNN, SVM, XGBoost, and RF again demonstrated strong performance at the 80:20 split, with peak accuracies of 86.7%, 80.0%, 80.0%, and 86.7%, respectively. QDA showed the steepest decline under data-scarce conditions, dropping to 36.4% accuracy at the 40:60 split. Across both classification tasks and all split ratios, CNN emerged as the most robust and reliable model, demonstrating resilience to data imbalance. These findings illustrate feasibility for detecting group-level spectral differences associated with tumor status using label-free SERS profiling of plasma sEVs.

### sEV proteomic profiling highlights molecular differences between tumor groups

To characterize the proteomic composition of sEVs in brain tumor patients, we performed LC-MS/MS analysis on proteins from plasma-derived sEVs (GB, *n* = 17; MNG, *n* = 20; HC, *n* = 30) from subjects of the same cohort analyzed by SERS. We identified and quantified sEV-associated proteins across all groups, revealing distinct and overlapping proteomic signatures (**Fig. 4**). A total of 421 proteins were identified in all three groups, while subsets of 431, 328, and 240 proteins were uniquely identified in HC, MNG, and GB groups, respectively (**Fig. 4a**). Although sEVs can cross the blood-brain barrier^91^, the majority of proteins detected in our dataset were plasma- and immune-related rather than brain- or tumor-specific, reflecting the strong influence of systemic circulation on sEV cargo profiles. These distributions potentially underscore both shared sEV biology and tumor-specific molecular features. Quantitative analysis of protein abundance revealed marked divergence in sEV protein profiles between GB and MNG samples (**Fig. 4b**). GB-derived sEVs were enriched for lipid transport and immune-modulatory proteins including APOC3, RBP4, and CLU, whereas MNG-derived sEVs showed higher levels of proteins such as IGLV2-14, LUM, and BTD.

**Figure 4.**
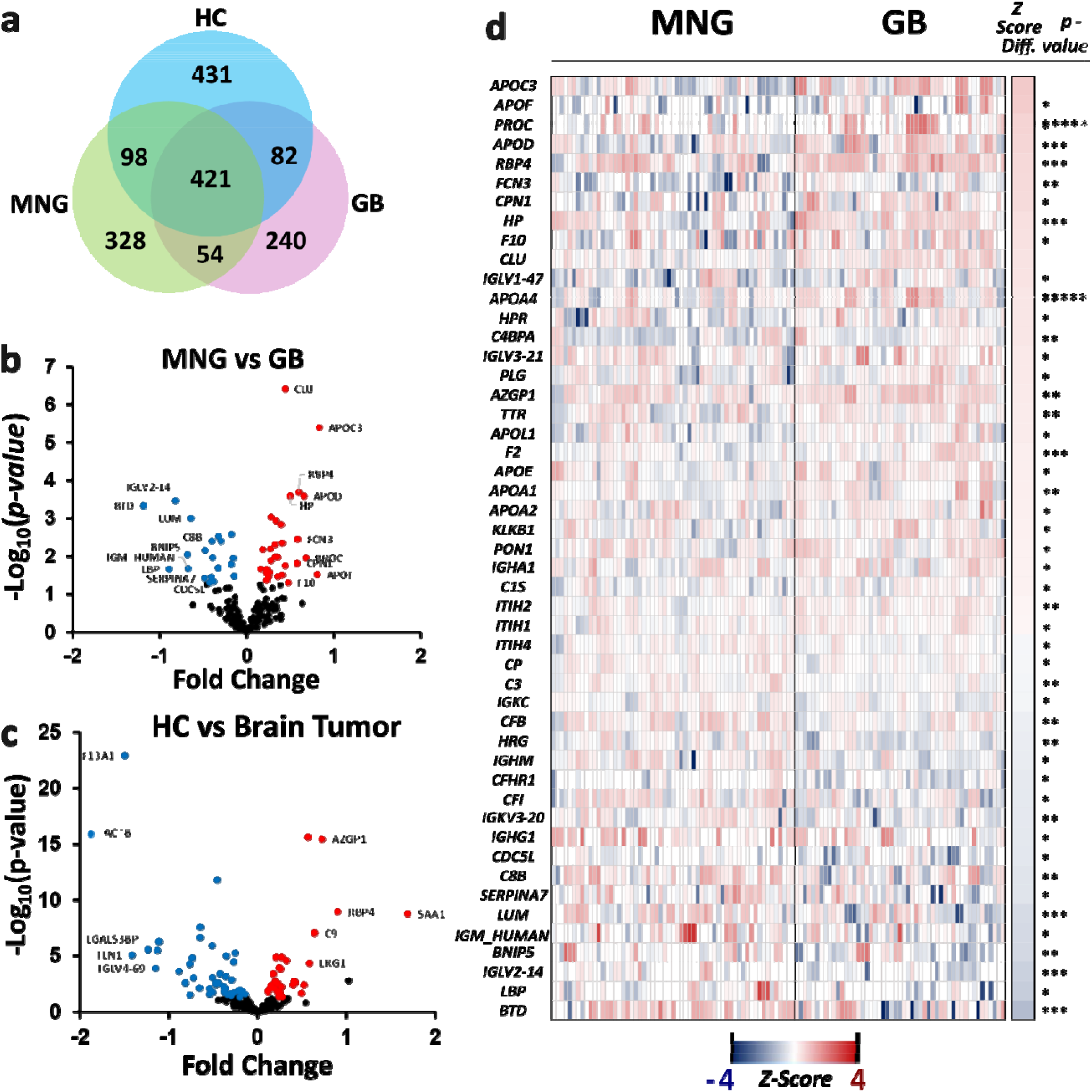
Proteomic Analysis of Plasma Small Extracellular Vesicles (sEVs) Reveals Differences in Protein Levels Between Sample Types. Proteomics analysis of plasma small extracellular vesicles (sEVs) derived from glioblastoma (GB), meningioma (MNG), and healthy control (HC) samples. (**a**) Venn diagram of identified proteins detected in at least one injection per sample across GB, MNG, and GB groups. The volcano plots (**b,c**) show the differential protein abundances between MNG vs GB and HC vs brain tumor (GB and MNG) groups. The x-axis represents the fold change (log_2_ scale) of protein abundance, while the y-axis shows the statistical significance as -log_10_(p-value). Proteins with a significant increase in abundance in one group are shown in red (positive fold change), while proteins significantly more abundant in the other group are displayed in blue (negative fold change). Non-significant proteins are shown in black. Volcano plot of differentially abundant proteins between (**b**) MNG vs GB and (**c**) HC vs brain tumor samples. (**d**) Heatmap of differentially abundant proteins (P < 0.05) between MNG and GB samples. The heatmap displays proteins with a significant difference in abundance (p < 0.05) when comparing MNG and GB groups. Rows represent individual proteins, while columns represent samples within each group. Z-scores indicate relative abundance levels, with red representing higher abundance and blue having lower abundance. The proteins are ranked based on their Z-score differences between groups and the statistical significance, is represented by the asterisks (p < 0.05, ** p < 0.01, *** p < 0.001, **** p < 0.0001).

When comparing healthy controls (HC) with the combined tumor group (MNG + GB), we observed distinct shifts in sEV cargo composition (**Fig. 4c**). Plasma-derived sEVs exhibited elevated levels of inflammation- and lipid-associated proteins, including A2GP1, SAA1, and LRG1. In contrast, HC-derived sEVs were enriched for structural and coagulation-related proteins such as ACTB, F13A1, and TNNI2. Hierarchical clustering further demonstrated separation of GB and MNG sEV profiles by tumor type (**Fig. 4d**). GB samples consistently displayed higher levels of proteins such as APOF, FCN3, and APOA4, while MNG samples showed enrichment of PROC and CPN1. Statistical analysis confirmed significant differences in levels of several proteins, with CLU and APOC3 among the most upregulated in GB (*****, *p* < 0.0001), and IGLV2-14 and LUM significantly enriched in MNG. These protein-level differences provide a parallel analysis to SERS-based sEV detection. Furthermore, protein-associated SERS bands, including phenylalanine (1002 cm□^1^), Amide I/II/III, and CH□/CH□ bending modes, showed group-associated intensity differences at the patient level across GB, MNG, and HC samples **(Supplementary Fig. 10)**.

## Discussion

The SERS spectral profiles of circulating sEVs can vary depending on physiological and pathological conditions^92^. Convolutional neural networks (CNNs) are particularly well suited for analyzing high-dimensional spectral data, including one-dimensional SERS inputs^93,94^. In our results, the CNN model achieved he strongest classification performance across evaluated models, with improved accuracy, specificity, and sensitivity supporting the feasibility of deep learning-based molecular pattern recognition for spectral analysis. Importantly, our results highlight the relevance of patient-wise data splitting as the primary evaluation strategy, as spectra-wise splitting may introduce overfitting due to shared spectral features between training and test sets. This distinction is critical for translational applications, where model performance must reflect patient-level generalizability rather than spectra-level similarity.

Proteomics analyses further support the potential of our sEV-based approach. In GB, elevated CLU^95,96,97^ and APOC3^98^ have been associated with immune modulation and metabolic processes^96,98^, while MNG-specific proteins such as IGLV2-14^99^, BTD, and LUM^100^ may reflect differences in tumor microenvironment interactions^100,101^. In addition to tumor-associated signals, systemic differences were observed between healthy controls and patients, including proteins related to coagulation^102,103^ (F13A1), cytoskeletal regulation (ACTB), metabolism and immune pathways (RBP4^104^, AZGP1^105,106^), and complement and angiogenesis (C9^107^, LRG1^108^). Together, these findings suggest that plasma sEV proteomics captures a combination of tumor-associated and systemic biological signals, providing complementary molecular context for interpreting SERS-derived spectral variation. This study presents a proof-of-concept evaluation of SERS-based molecular characterization of patient-derived plasma sEVs. Our results demonstrate the feasibility of integrating label-free spectroscopy with AI to identify group-associated molecular differences between patient cohorts.

The two-stage framework explored here, detection of brain tumor-associated signals followed by exploratory tumor-type stratification, provides a conceptual basis for future investigation, but should be interpreted cautiously given the pilot cohort size. Given the molecular heterogeneity of high-grade gliomas, validation in larger and clinically representative cohorts, including relevant control populations and additional tumor subtypes, will be required to assess robustness and generalizability. Expanding the dataset will also enable improved model training and evaluation of whether more subtle spectral differences can be consistently detected across heterogeneous tumor populations. Standardization of sample collection and processing will be particularly important, as plasma samples from GB and MNG patients were collected from patients prior to tumor resection, whereas healthy control samples were obtained from walk-in donors. Plasma-derived sEVs represent a heterogeneous mixture of vesicles originating from multiple systemic sources, making it difficult to isolate tumor-specific signals.

In conclusion, this study demonstrates the successful isolation and characterization of plasma-derived sEVs integrated with label-free SERS analysis and AI-based classification. Using clinical samples, as a proof of concept, our findings show that this combined approach can capture group-associated molecular differences between sample cohorts. These results should be considered preliminary and primarily methodological, supporting the feasibility of combining EV isolation, SERS, and AI for molecular profiling of plasma samples. Further studies will be needed to assess model performance across a broader range of brain tumor types and clinical contexts.

## Acknowledgments

The Canary Foundation, the Beckman Center for Molecular and Genetics Medicine, and the NIH R25 CREST program are acknowledged by the authors for their support. The authors acknowledge the use of the services and facilities of the Koç University Hospital Clinical Trials Unit and Koç University Research Center for Translational Medicine (KUTTAM). U.A. acknowledges funding from the European Union’s Horizon Europe research and innovation program under the Marie Skłodowska-Curie grant agreement No. 101066038. We greatly acknowledge Drs. Sharon Pitteri, Fernando Jose Garcia Marques, and Abel Bermudez for their support in the proteomics analysis. Drs. Ramasamy Paulmurugan and Arutselvan Natarajan are acknowledged for their support in flow cytometry experiments. Dr. Torun presented the initial version of this work at ISEV 2024 and gratefully acknowledges the ISEV 2024 Research Scholarship.

## Funding

**N/A**

## Author contributions

H.T., U.P. and U.D. have full access to all data and take responsibility for the integrity and accuracy of the data analysis; Concept and design: H.T., I.S., U.D.; Acquisition, analysis, or interpretation of data: H.T., U.P., C.N., T.V., F.K., O.A., U.A., B.S., I.K., D.A., I.S., U.D.; Drafting of the manuscript: H.T., C.N.; Critical revision of the manuscript for important intellectual content: H.T., U.P., D.A., U.A., M.O.O., I.S., U.D.; Statistical analysis: H.T., U.P., T.V.; Obtained funding: I.S., U.D.; Administrative, technical, or material: H.T., U.P., C.N., T.V., O.A., F.K., R.C.E., I.K., G.A., O.B., I.S., U.D.; Supervision: I.S., U.D.

## Competing interests

Utkan Demirci (UD) is a founder of and has an equity interest in: (i) Levitas Inc., (ii) Hermes Biosciences, (iii) Vetmotl Inc. UD’s interests were viewed and managed in accordance with the conflict-of-interest policies. All other authors declare they have no competing interests.

## Data and materials availability

The main data supporting the results in this study are available within the paper and its Supplementary Information. The raw and analyzed datasets generated during the study are too large to be publicly shared, but they are available for research purposes from the corresponding author on reasonable request. The patient pathology data are available from the authors, subject to approval from the Koç University Institutional Review Board.

