## Supplementary material for "Glio-SERS: Label-Free Molecular Profiling of Plasma Extracellular Vesicles in Brain Tumors Using SERS and Artificial Intelligence": GlioSERS_supplementary_materials_03182026.docx

Hulya Torun *et al.*

**This PDF file includes:**

Supplementary Text

Figs. S1 to S10

Tables S1 to S2

**Other Supplementary Materials for this manuscript include the following:**

Movies S1 to S1

Supplementary Text

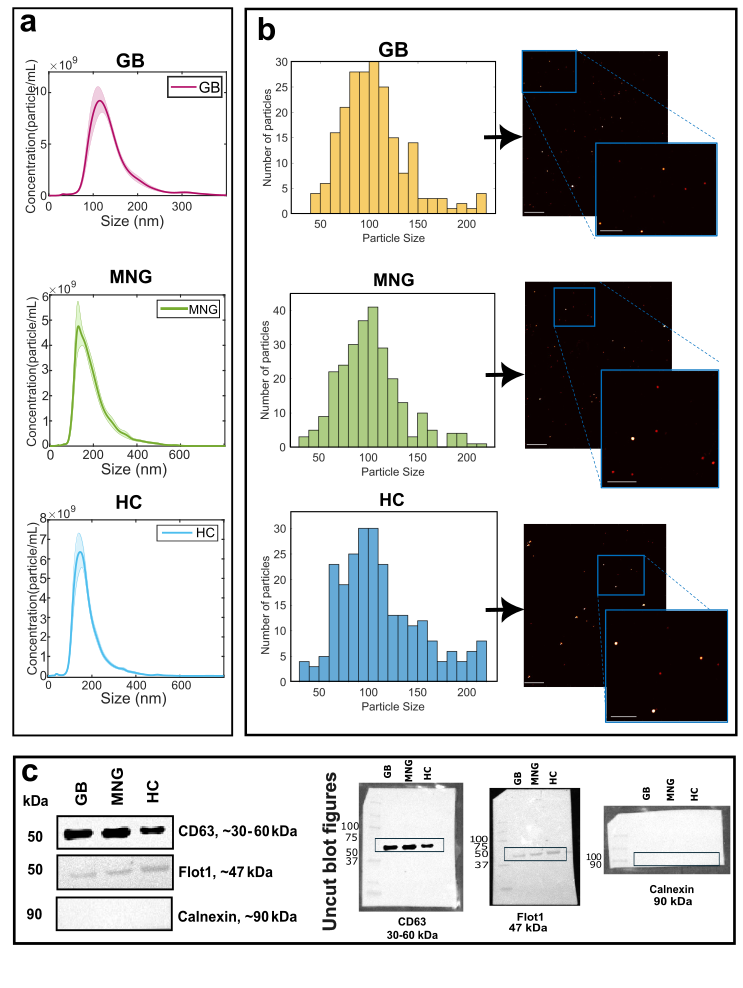

Fig. S1. Characterization of small extracellular vesicles derived from glioblastoma, meningioma and healthy control samples. sEVs were characterized by using a, Nanoparticle Tracking Analysis (NTA), b, Interferometric Scattering Light Microscopy (iSCAT), and c, Western blotting using CD63, Flot1, and Calnexin antibodies (both cropped and uncut figures provided on left and right side, respectively).

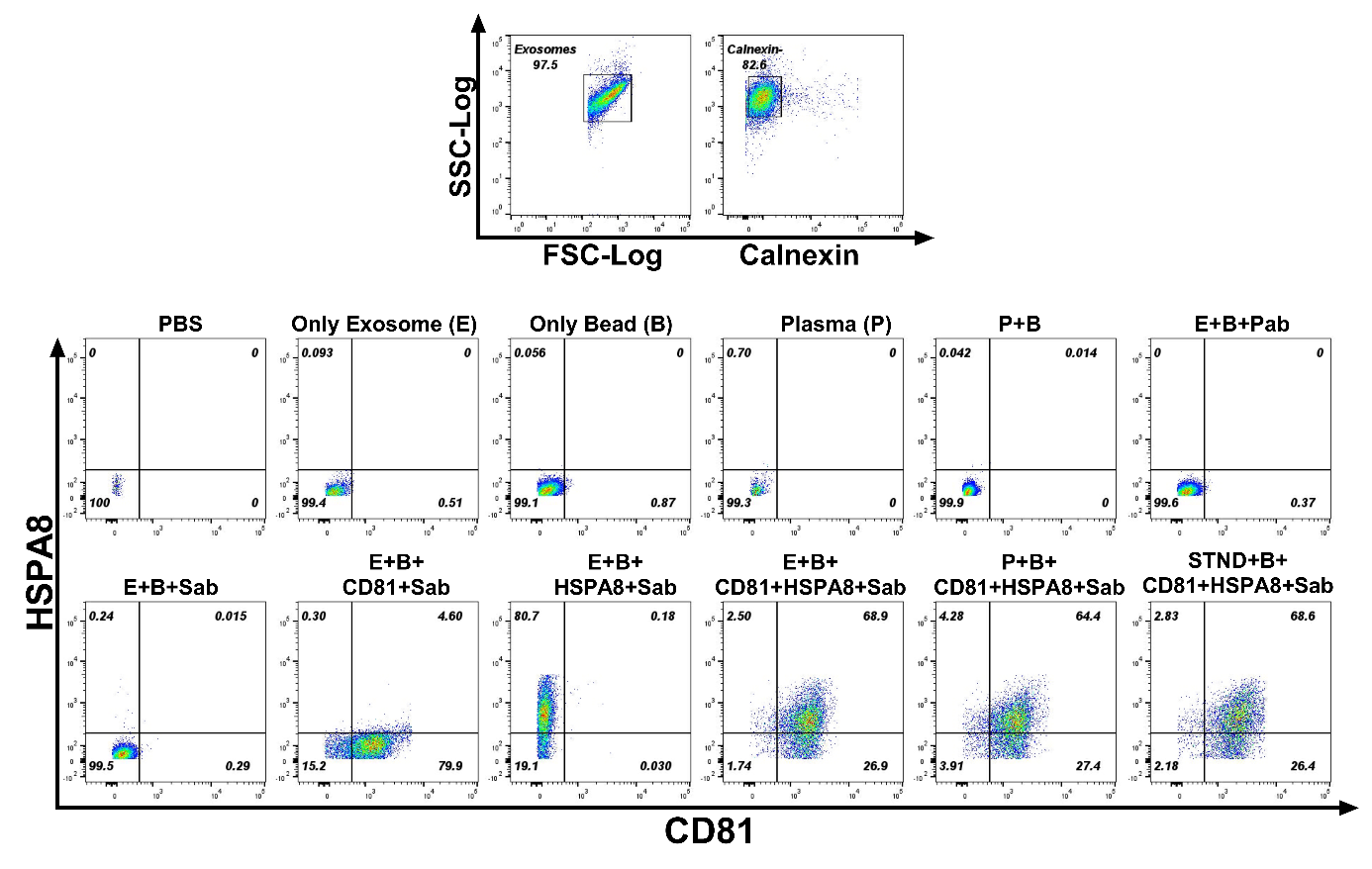

Fig. S2. Flow cytometry characterization of small extracellular vesicles (sEVs). sEVs were isolated and analyzed by flow cytometry following capture on aldehyde/sulfate latex beads. Representative gating strategy showing FSC-Log vs. SSC-Log profiles of sEVs (left), and staining with calnexin, an endoplasmic reticulum marker used as a negative control to confirm the absence of cellular contamination (right). (Bottom) Pseudocolor plots display the expression of sEV surface markers CD81 and HSPA8 under various experimental conditions: PBS (negative control), only sEVs (E), only beads (B), plasma alone (P), plasma plus beads (P+B), sEVs plus beads plus primary antibody (E+B+Pab), sEVs plus beads plus secondary antibody only (E+B+Sab), single-marker staining for CD81 (E+B+CD81+Sab) or HSPA8 (E+B+HSPA8+Sab), dual-marker staining for CD81 and HSPA8 (E+B+CD81+HSPA8+Sab), plasma-derived sEVs stained for CD81 and HSPA8 (P+B+CD81+HSPA8+Sab), and standard exosome preparations stained for CD81 and HSPA8 (STND+B+CD81+HSPA8+Sab). The use of CD81, a tetraspanin commonly enriched in sEVs, and HSPA8, a heat shock protein associated with EV biogenesis, enables phenotypic confirmation of sEV identity. Appropriate negative and FMO controls verify assay specificity and rule out non-specific bead or antibody binding.

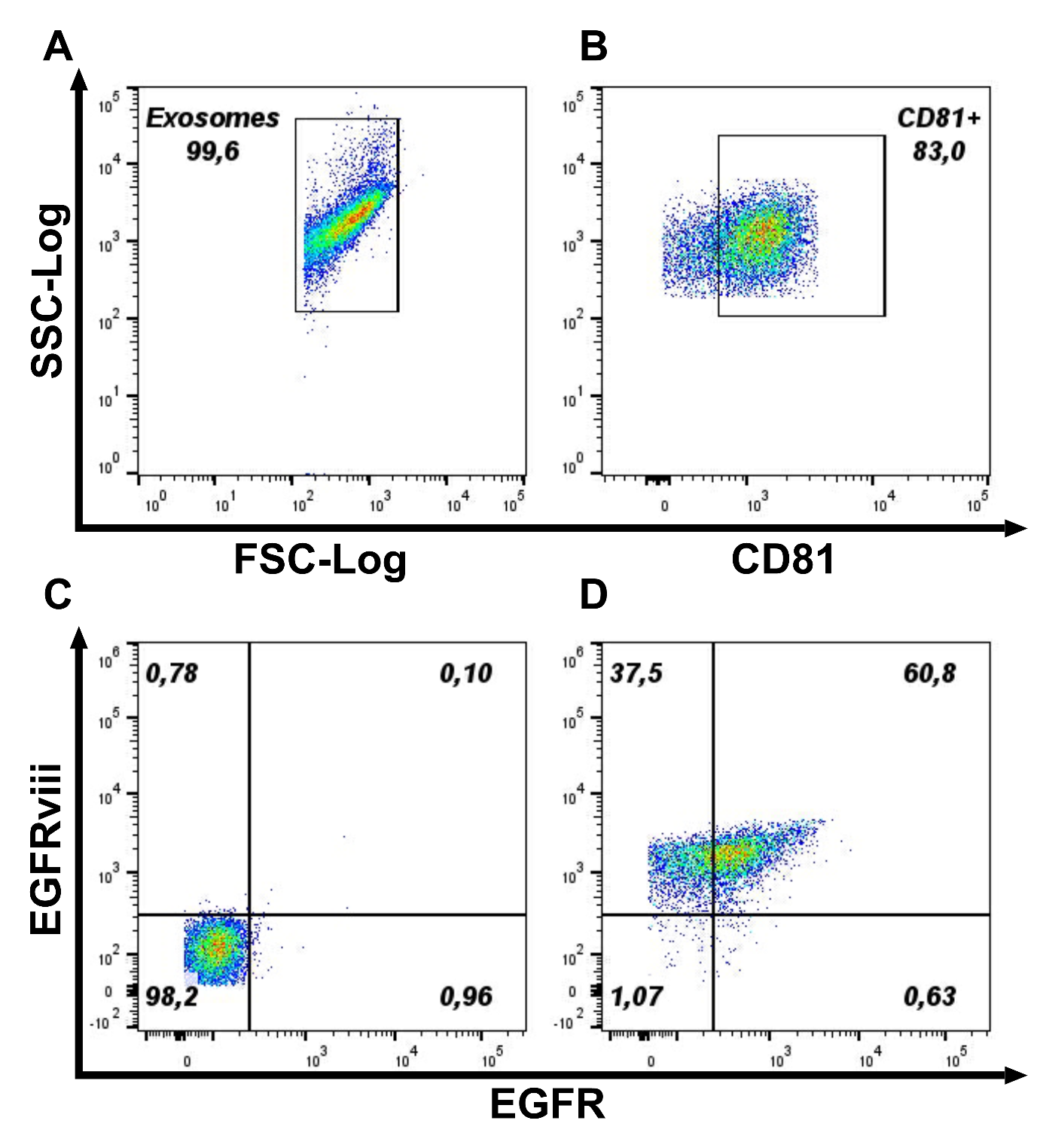
Fig. S3. Representative expression analysis of EFGR+ and EFGRviii+ on CD81+ small extracellular vesicles (sEVs). LogFSC and LogSSC scales were used to determine small scale particles. a, Events were gated from the FSCxSSC plot. b, sEVs were determined as CD81+ events. c, A sample unstained for EFGR/EFGRviii were used to gate out negative population. d, Representative analysis for EFGR+ and EFGRviii+ sEVs.

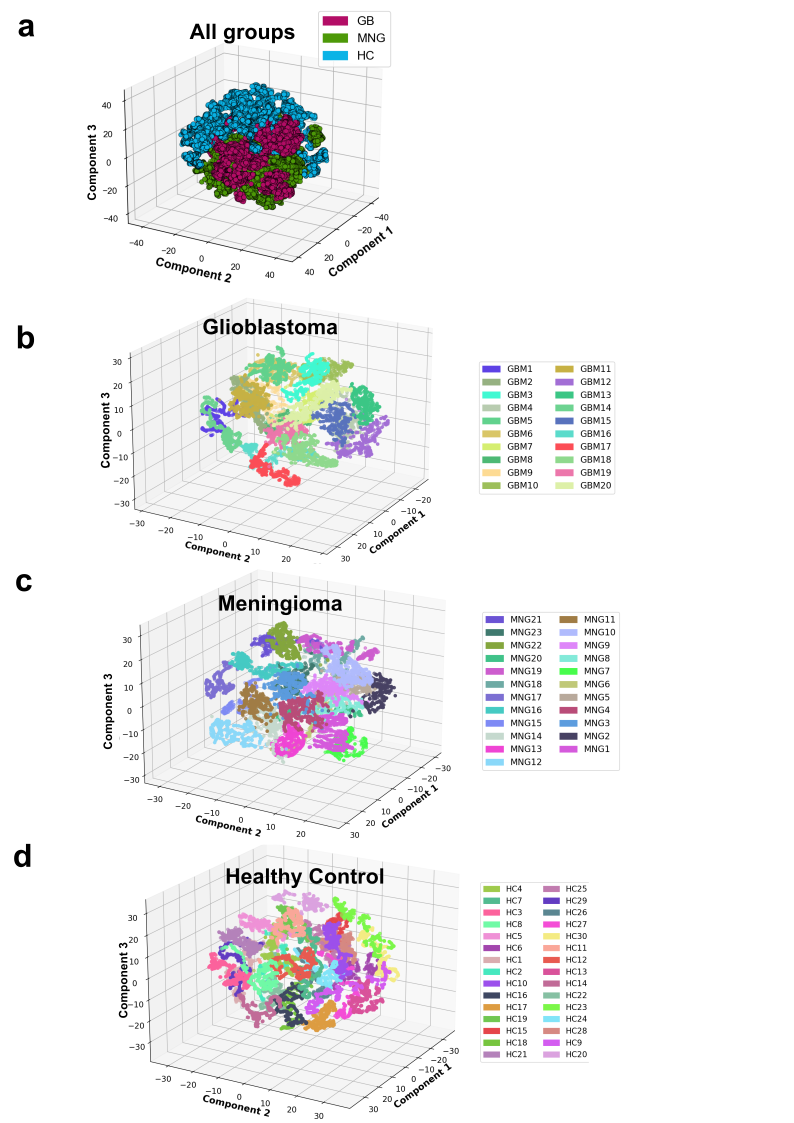

Fig. S4. t-SNE distribution of SERS spectra for a, all three groups, b, glioblastoma, c, meningioma, and d, healthy control EVs.

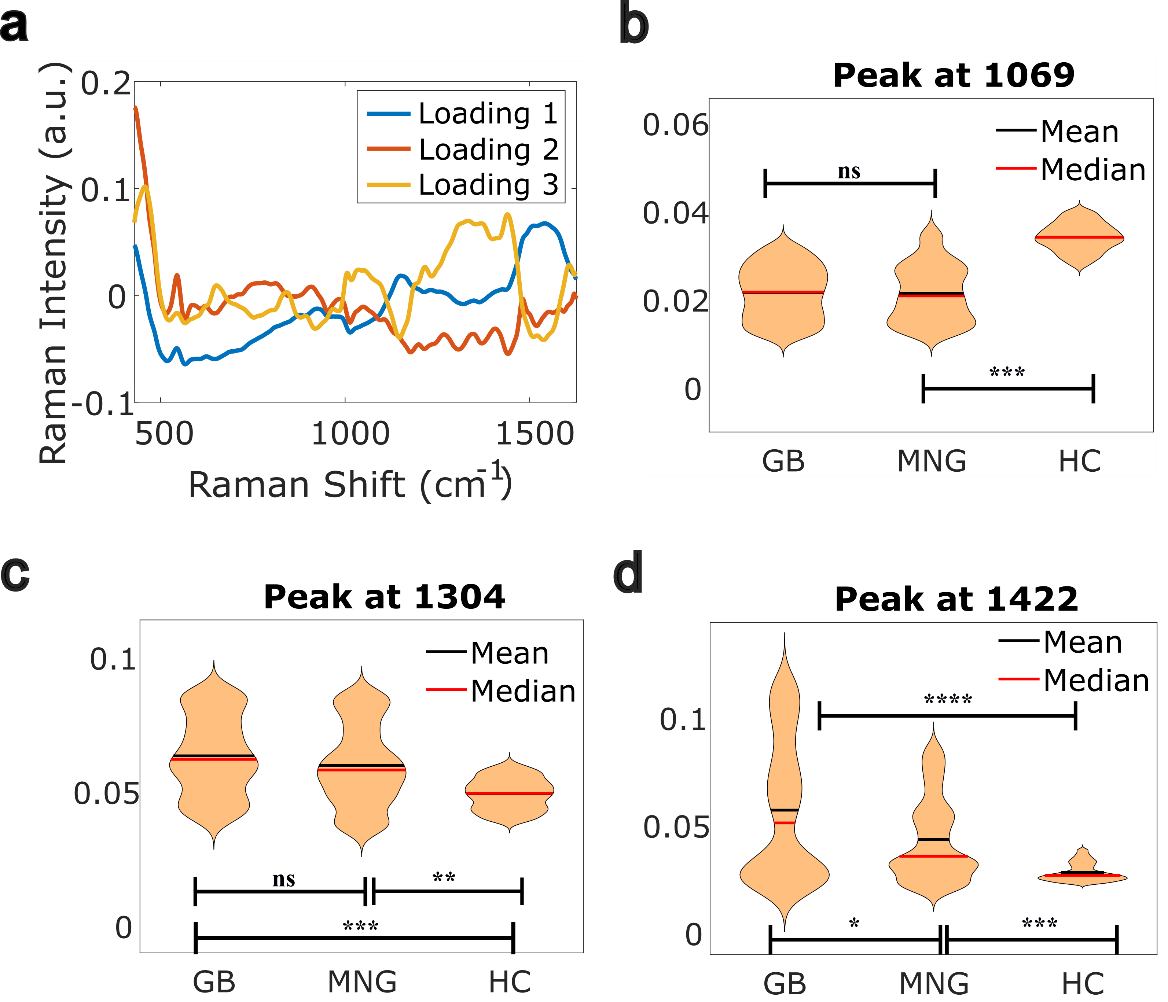

**Fig. S5.** PCA coefficients and **a**, violin plots for distinguishing Raman shifts in between 3 groups: **b**, glioblastoma, **c**, meningioma, and **d**, healthy control EVs. Statistical significance shown as p < 0.05 (*), < 0.01 (**), < 0.001 (***), < 0.0001 (****), n.s. not significant.

**AI Analysis of Extracellular Vesicle SERS signatures**

**Reproducibility and Random Seed Selection**

To ensure reproducibility and transparency of our computational analyses, random seeds were fixed throughout data preprocessing, spectra selection, and neural network initialization. Specifically, the train-test splits (both patient-wise and spectra-wise) and initialization parameters of the neural network (NN) and convolutional neural network (CNN) models were determined by predefined random seeds, set once by the authors. These seeds guarantee identical outcomes whenever the computational notebook is executed, independent of user, location, or timing of execution. It is important to emphasize that altering the random seeds may influence model performance and thus slightly affect the reproducibility of the results presented.

**Dropout Layers and Stability of CNN Predictions**

We observed minor variability in predictive accuracy across repeated evaluations of our saved CNN model. This variability arises from the use of dropout layers within the CNN architecture, which introduces a stochastic element by randomly deactivating a fraction of neurons in hidden layers. Dropout is commonly employed to reduce model overfitting and enhance generalizability across diverse datasets. Empirical evidence obtained in our analysis indicated that incorporating dropout substantially improved the robustness and generalization performance of the CNN classifier. Moreover, prior studies consistently support the effectiveness of dropout in improving the performance of neural networks across various image-based datasets. Given that SERS spectral vectors essentially represent one-dimensional image data, employing dropout layers in our CNN model was both appropriate and advantageous for enhancing predictive robustness and overall generalizability.

**EGFR Quantification Analysis**

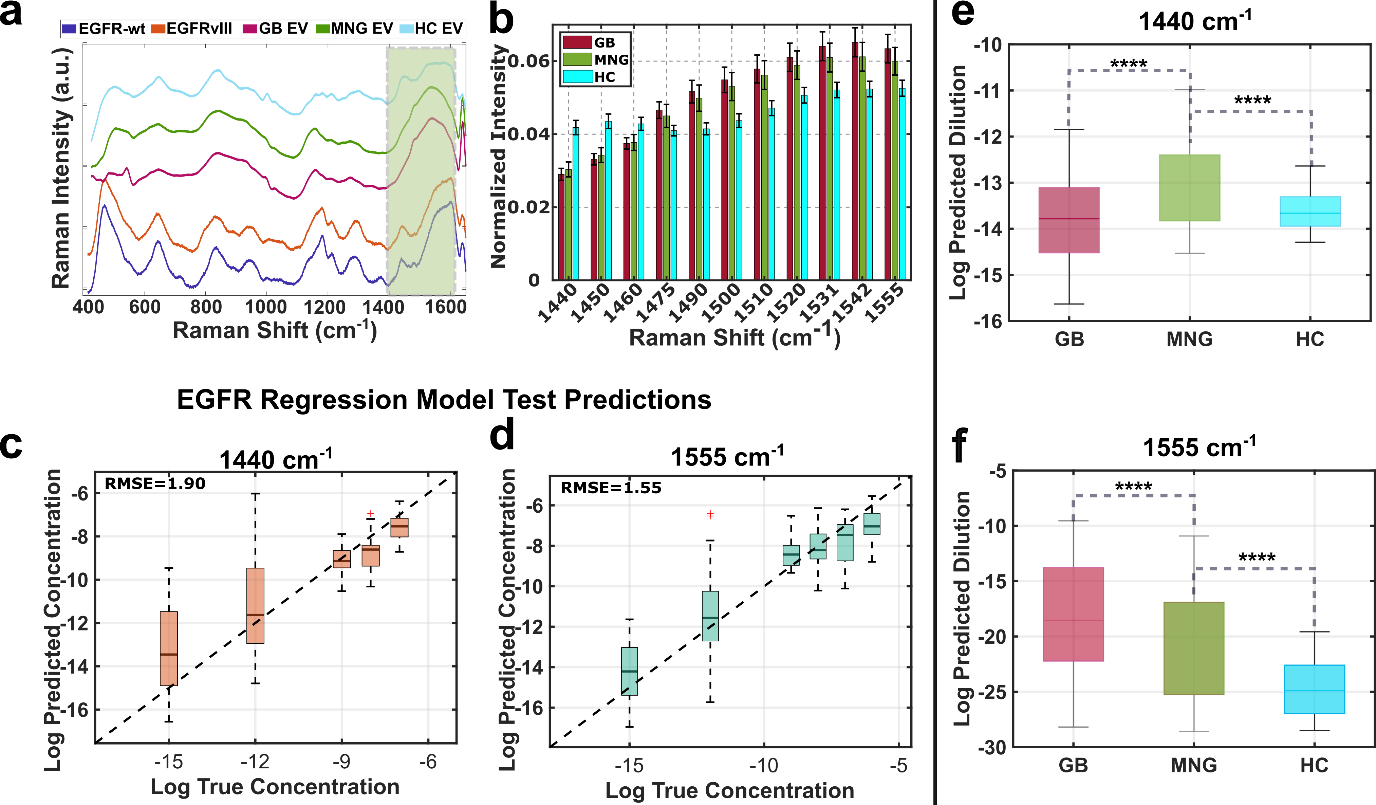
**Fig. S6. Raman-based detection and dilution quantification of EV-associated hepatotoxicity markers**. **a**, Representative Raman spectra from small extracellular vesicle (sEV) samples corresponding to three experimental groups: GB (orange), MNG (purple), and HC (blue). The shaded region (1440–1600 cm⁻¹) highlights significant spectral differences among the groups. **b**, Normalized mean intensities at Raman shifts in the 1440–1600 cm⁻¹ region, which were selected by performing statistical analysis on sEV and EGFR Raman spectra. Box plots of predicted vs. true log concentrations at **c**, 1440 cm^-1^, and **d**, 1555 cm^-1^. **e-f**, Log predicted concentration values from the trained regression model for each group, showing significantly lower predicted values for HC samples for models developed with the intensities at **e**, 1440 cm^-1^ and **f**, 1550 cm^-1^. Statistical significance shown as p < 0.05 (*), < 0.01 (**), < 0.001 (***), < 0.0001 (****), n.s. not significant.

| **Patient** | **Molecular Pathology (c/mL)** | **Raman Prediction (c/mL)** |
| --- | --- | --- |
| 5 | 13 | 4.3 |
| 7 | 12 | 3.4 |
| 10 | 8 | 3.5 |
| 12 | 4.5 | 5.2 |

Table S1. Comparison of predicted EGFR quantification from molecular pathology (Next Generation Sequencing, NGS) and developed SERS regression model.

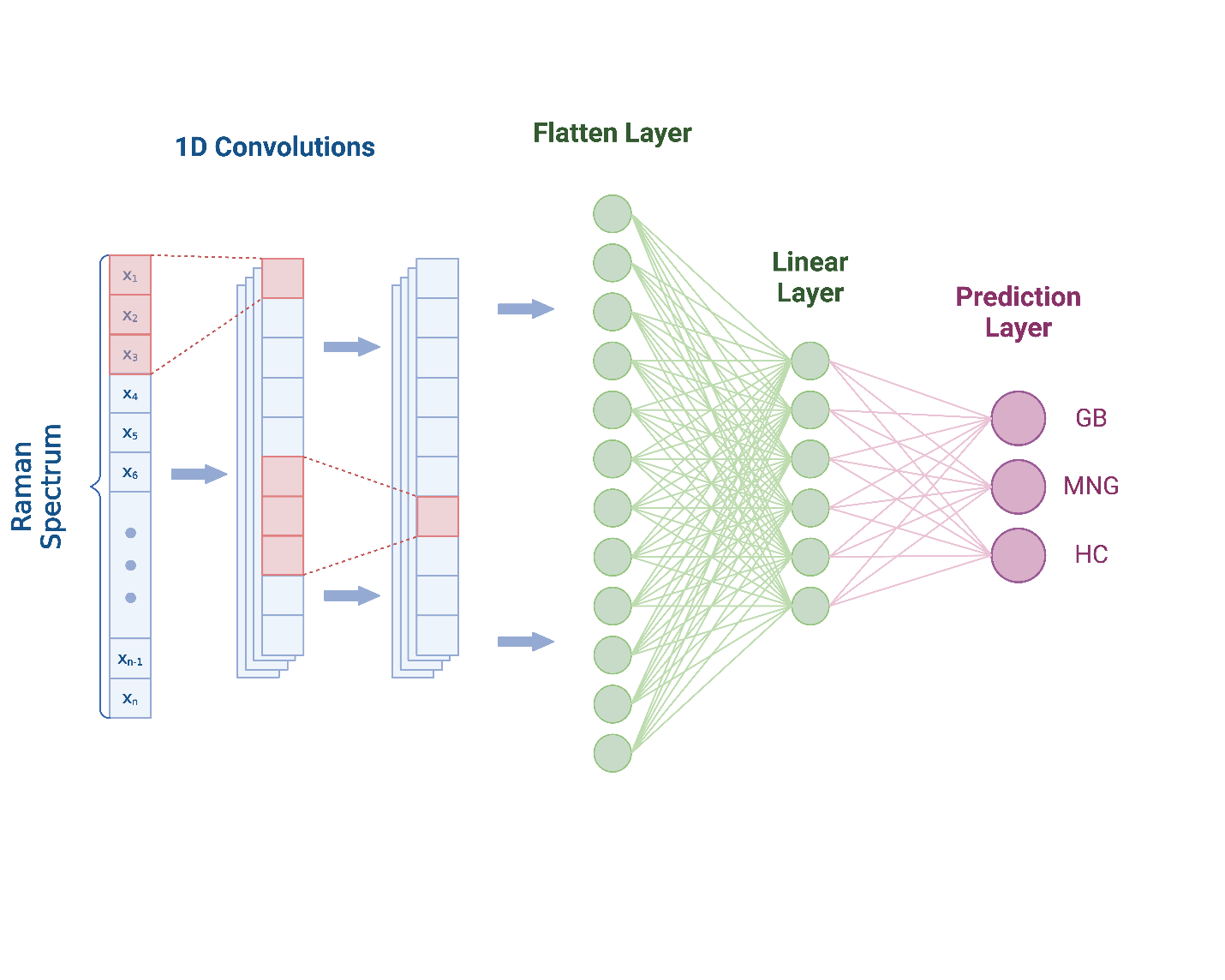

Fig. S7. Convolutional Neural Network (CNN) Architecture. The schematic representation of CNN model architecture.

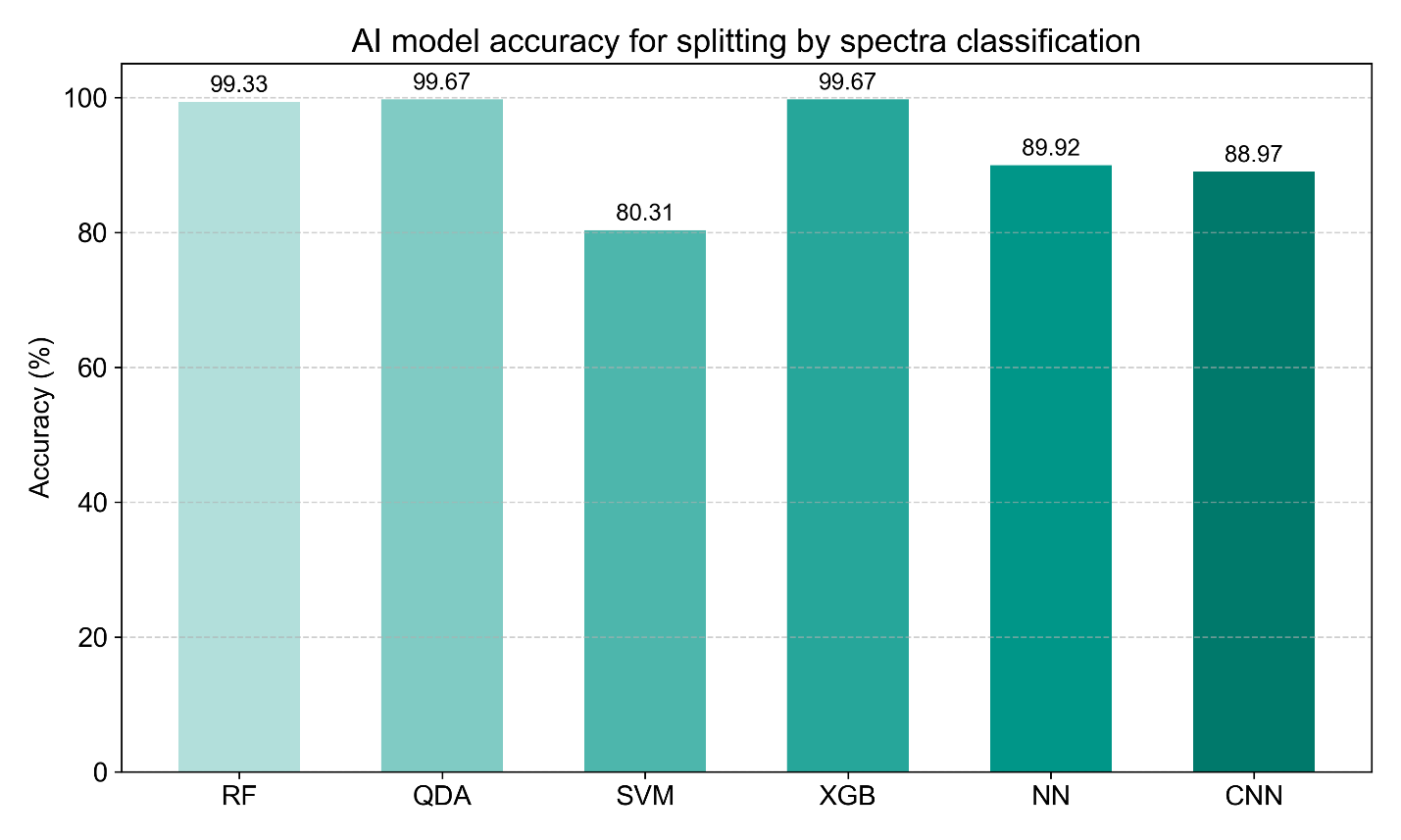

Fig. S8. Accuracy comparison for different AI models by using split by spectra classification method in classification of brain tumor (BT: GB and MNG) vs healthy control (HC) groups. (CNN: Convolutional Neural Networks, NN: Neural Networks, RF: Random Forest, SVM: Support Vector Machine, XGB: XGBoost, QDA: Quadratic Discriminant Analysis)

| **Models/ Train-Test Split Ratio** | **AUC 60:40** | **Accuracy** | **Specificity (BT vs HC)  for 60:40** | **Sensitivity (BT vs HC)  for 60:40** | **Weighted f1 Score  for 60:40** |
| --- | --- | --- | --- | --- | --- |
| **CNN** | 0.85 | 0.73 | 0.86 | 0.1 | 0.72 |
| **NN** | 0.85 | 0.26 | 0.66 | 0.31 | 0.18 |
| **RF** | 0.9 | 0.72 | 0.87 | 0.69 | 0.71 |
| **SVM** | 0.9 | 0.71 | 0.87 | 0.69 | 0.7 |
| **XGBoost** | 0.89 | 0.71 | 0.87 | 0.68 | 0.7 |
| **QDA** | 0.76 | 0.58 | 0.79 | 0.57 | 0.59 |
| **Models/ Train-Test Split Ratio** | **AUC 40:60** | **Accuracy** | **Specificity (BT vs HC)  for 40:60** | **Sensitivity (BT vs HC)  for 40:60** | **Weighted f1 score  for 40:60** |
| **CNN** | 0.79 | 0.67 | 0.83 | 0.64 | 0.66 |
| **NN** | 0.74 | 0.31 | 0.67 | 0.34 | 0.23 |
| **RF** | 0.86 | 0.67 | 0.85 | 0.65 | 0.66 |
| **SVM** | 0.9 | 0.69 | 0.86 | 0.65 | 0.66 |
| **XGBoost** | 0.85 | 0.65 | 0.83 | 0.63 | 0.64 |
| **QDA** | 0.58 | 0.38 | 0.69 | 0.46 | 0.39 |

**Table S2.** **AI classification results for brain tumor (BT: GB and MNG) vs healthy control (HC) groups are given for different train-test split ratios**: Area Under the Curve (AUC), Specificity, Sensitivity, and Weighted f1 Score values are presented for 60 train-40 tests ratio shown in pink, 40 train-60 test ratios shown in green.

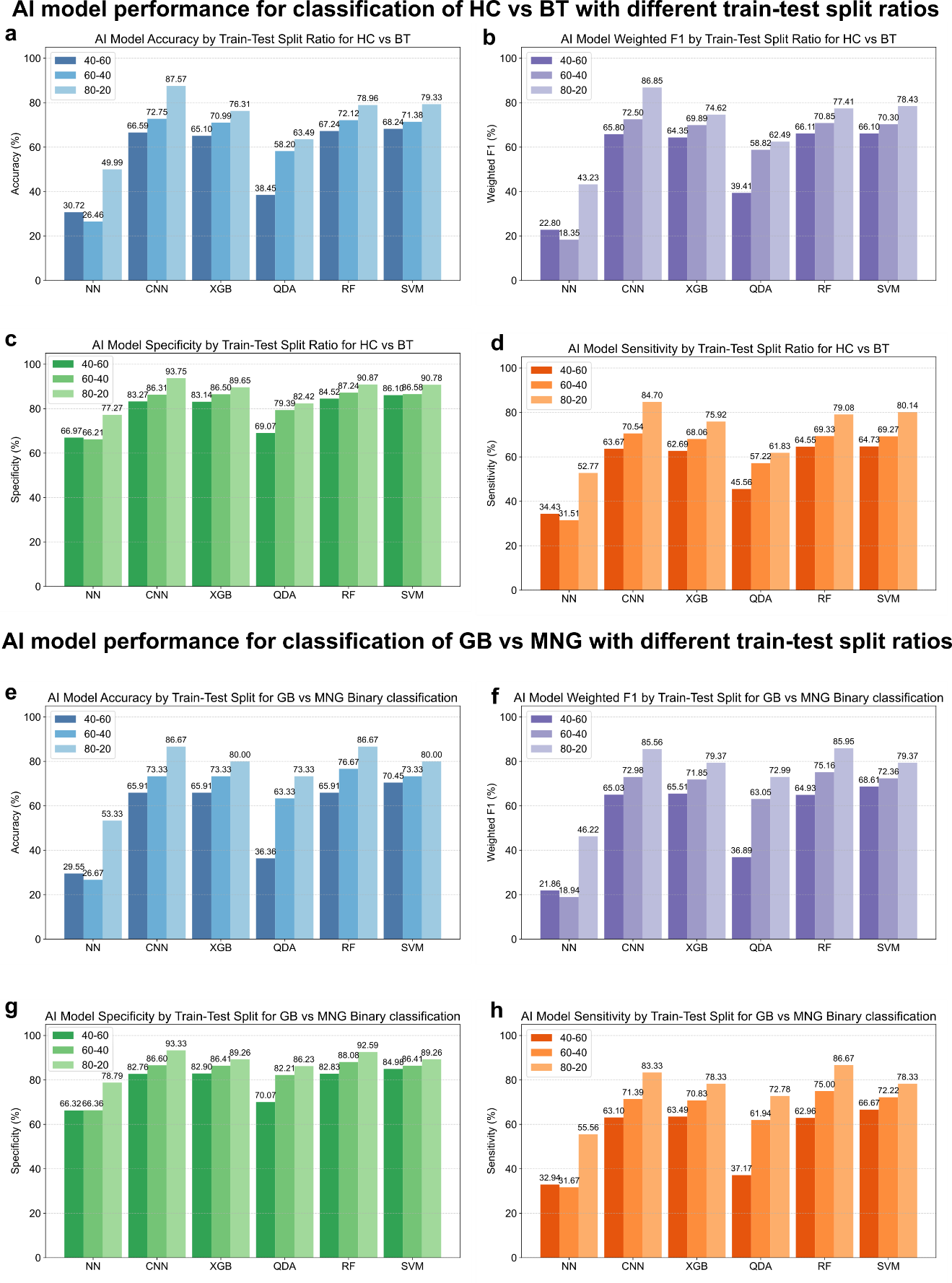

Fig. S9. Performance of artificial intelligence models for SERS-based classification of brain tumor subtypes and healthy controls under varying train-test split ratios using patient-wise split method. (CNN: Convolutional Neural Networks, NN: Neural Networks, RF: Random Forest, SVM: Support Vector Machine, XGB: XGBoost, QDA: Quadratic Discriminant Analysis). The diagnostic performance of multiple machine learning and deep learning models-including neural networks, convolutional neural networks (CNN), XGBoost (XGB), support vector machines (SVM), random forest (RF), and quadratic discriminant analysis (QDA)-was evaluated across different train-test split ratios for two classification tasks: healthy controls (HC) vs brain tumors (BT), and glioblastoma (GB) vs meningioma (MNG). a, Model accuracy across train–test split ratios for HC vs BT classification. b, Weighted F1 scores for HC vs BT classification. c, Specificity of each model across train-test split ratios for GB vs MNG classification. d, Sensitivity of each model for HC vs BT classification. e, Accuracy across models for GB vs MNG binary classification. f, Weighted F1 scores for GB vs MNG classification. g, Specificity of models for GB vs MNG classification. h, Sensitivity of models for GB vs MNG classification. Each point reflects performance metrics computed on the held-out test set, with models trained and evaluated independently under the indicated data splits.

SERS protein-related features in sEV spectra across groups

To interrogate protein-linked differences among GBM, MNG, and HC sEVs, we scanned the calibrated Raman axis and then focused analysis on protein-dominated bands used in our code: 1002 cm^-1^ (phenylalanine ring breathing); Amide III windows 1245–1265, 1270–1295, and 1300–1315 cm^-1^; CH₂/CH₃ bending 1442–1460 cm^-1^; Amide II 1540–1565 cm^-1^; aromatic ring 1605–1615 cm^-1^; and Amide I 1645–1665 cm^-1^.

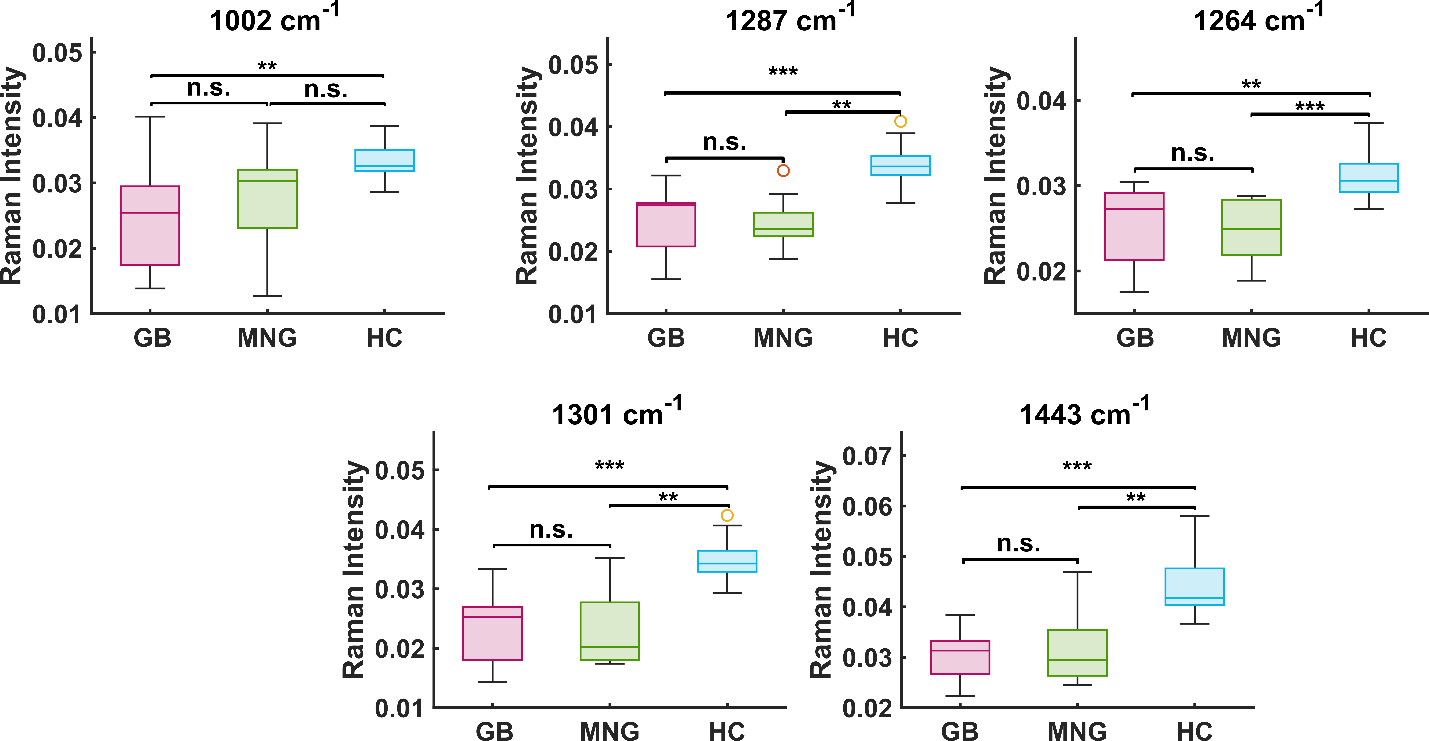

Fig. S10. | Protein-related Raman bands at 1002, 1255, 1304, 1450, 1610, 1640 cm^-1^ are compared in GB, MNG, and HC groups. Pairwise differences were tested with two-sided Mann–Whitney U tests and FDR (BH) across all bands. Statistical significance shown as p < 0.05 (*), < 0.01 (**), < 0.001 (***), < 0.0001 (****), n.s. not significant.

Analysis was performed at the patient level to avoid pseudo replication: each patient’s spectra was averaged, robust outliers were removed (Tukey 1.5×IQR (Interquartile range) with an optional MAD=3 gate), and pairwise group differences at every wavenumber were tested using two-sided Mann-Whitney U. We controlled multiplicity across all wavenumbers × group pairs with Benjamini-Hochberg FDR and quantified effects with Cliff’s delta (δ). Among the protein-related windows, we retained positions with q < 0.05 and ranked them by a composite score S = weight × |δ| × (−log_10_ q) (weights reflect prior confidence in protein specificity). The top-ranked bands (e.g., Amide I 1645–1665 cm^-1^, CH₂/CH₃ bending 1442-1460 cm^-1^, and phenylalanine 1002 cm^-1^) are displayed as patient-level box plots with FDR-adjusted significance brackets. Across these bands, GB samples frequently showed elevated intensities relative to MNG and/or HC, consistent with enhanced protein/lipid signatures in tumor-derived vesicles.

Movie S1. 3D visualization of PCA components.

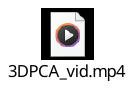
